# A bivalent Lassa virus mRNA vaccine protects guinea pigs from lethal infection with Lassa virus

**DOI:** 10.64898/2026.10.25.761704

**Authors:** Mei Hashizume, Takeshi Saito, Ryotaro Nouda, Takashi Matsuzaki, Shintaro Yamada, Gavriella Siman-Tov, Daiki Kobayashi, Christine Click, Kirsten Littlefield, Emily Schugardt, Igor S. Lukashevich, Juan C. de la Torre, Junki Maruyama, Masaharu Iwasaki

**Author notes:** Corresponding authors. (J. Maruyama), (M. Iwasaki).

## Abstract

Lassa virus (LASV), the causative agent of the life-threatening febrile disease, Lassa fever, remains a major public health concern in West Africa, where hundreds of thousands of infections occur annually. However, no licensed vaccine is currently available, highlighting the urgent need for effective preventive measures. We previously demonstrated that an mRNA-lipid nanoparticle (LNP) vaccine expressing LASV glycoprotein precursor (GPC; LASgpc-mRNA-LNP) conferred protection against lethal infection with a recombinant lymphocytic choriomeningitis virus expressing LASV GPC in mice. In this study, we extended our previous work with the aim of evaluating the efficacy of an mRNA-LNP vaccine expressing LASV nucleoprotein (NP; LASnp-mRNA-LNP) and a bivalent vaccine combining LASgpc-mRNA-LNP and LASnp-mRNA-LNP. Using a lethal CBA/N mouse model of infection with ML29, an LASV live-attenuated vaccine candidate encoding LASV GPC and NP, we found that LASgpc-mRNA-LNP and LASnp-mRNA-LNP each conferred protection in the absence of detectable neutralizing antibody responses. We identified LASV GPC- and NP-derived CD4⁺ and CD8⁺ T-cell epitope peptides in CBA/N mice and demonstrated that vaccination induced robust antigen-specific T-cell responses. Importantly, the bivalent vaccine provided complete protection against lethal challenge with doses at which monovalent vaccines failed to protect all mice, suggesting a potential dose-sparing benefit of the bivalent formulation. Furthermore, the bivalent vaccine fully protected guinea pigs against lethal challenge with authentic LASV. Together, these findings establish LASV NP, in addition to GPC, as a protective antigen in the mRNA vaccine platform and provide a rationale for developing bivalent mRNA-LNP vaccines to improve protection against Lassa fever.

**One Sentence Summary:** A bivalent mRNA-LNP vaccine expressing Lassa virus GPC and NP confers robust protection against lethal Lassa virus infection

## INTRODUCTION

Lassa virus (LASV), a member of the family *Arenaviridae* and the genus *Mammarenavirus*, is the causative agent of Lassa fever (LF), a life-threatening acute febrile disease endemic to large areas of West Africa (*1, 2*). Recent modeling studies have estimated that LASV infects 2.7 million individuals each year, resulting in more than 24,000 cases requiring hospitalization (*3*). LASV Enable 1.0 Research (2020–2023) across 113 rural communities in Benin, Liberia, Nigeria, and Sierra Leone documented that children have a twice higher risk of contracting LF in West Africa than adults (*4*). Case fatality rates (CFRs) among hospitalized, laboratory-confirmed LF cases vary considerably across regions and outbreak clusters, with an exceptionally high CFR of 60% reported during the 2015–2016 outbreak in Nigeria (*5*). Even with enhanced disease surveillance and response capabilities, Nigeria reported a CFR of 19% in 2025 (*6, 7*). LF was designated a World Health Organization (WHO) R&D Blueprint Priority Disease in 2016, and it remained on the updated priority list in 2018 (*8*). Despite its significant public health impact, there are currently no licensed vaccines for LF, and limited therapeutic options involve the off-label use of the nucleotide analog ribavirin, which is partially efficacious and has significant side effects (*9*). Because limited access to medical facilities and healthcare-seeking behavior can pose challenges to timely treatment in some settings, vaccination is considered a promising strategy for reducing the burden of LF in endemic regions (*10, 11*). Accordingly, LASV has been designated a priority pathogen by the Coalition for Epidemic Preparedness Innovations, which has supported the development of multiple LASV vaccine candidates, with some advancing to clinical trials (*12*).

Early activation of cellular immune responses is widely recognized as a critical determinant for controlling LASV infection in LF survivors and experimental animal models (*13–16*). Additionally, the limited healthcare infrastructure in some LASV-endemic regions highlights the need for vaccines that induce durable protective immunity following a single administration, as emphasized in the WHO Target Product Profile for Lassa virus Vaccine (*17*). Live-attenuated vaccines (LAVs) are well suited to these requirements because they induce robust, long-lasting cellular immunity after a single dose. Among various LASV LAV candidates (*18*), rVSVΔG-LASV-GPC, a recombinant vesicular stomatitis virus (VSV)-based vaccine expressing the LASV glycoprotein precursor (GPC) in place of the native VSV glycoprotein, is currently the most clinically advanced candidate, having completed phase 1 evaluation and progressed to an ongoing phase 2 trial (NCT05868733) (*19*). Nevertheless, safety concerns associated with live viral-vectored vaccines remain, particularly regarding their use in populations vulnerable to LF, including immunocompromised individuals, children, and pregnant women (*20*). mRNA-lipid nanoparticle (LNP) vaccines have emerged as a powerful and scalable platform, as demonstrated by the rapid development and global deployment of coronavirus disease 2019 (COVID-19) vaccines encoding the severe acute respiratory syndrome coronavirus 2 spike protein (SARS2-S) (*21–23*). Unlike live viral-vectored vaccines, mRNA vaccines contain no infectious agents and may therefore mitigate some of the safety concerns associated with LAVs. Importantly, studies in humans and animals have shown that COVID-19 mRNA-LNP vaccines induce neutralizing antibodies while eliciting SARS2-S-specific T-cell responses, supporting the potential application of this technology to LASV vaccine development, for which cellular immunity is considered critical (*24, 25*). We and others demonstrated that mRNA-LNP vaccines expressing LASV GPC (LASgpc-mRNA-LNP) conferred strong protective immunity in CBA/N mice challenged with recombinant lymphocytic choriomeningitis virus (LCMV), a related mammarenavirus, expressing LASV GPC (rLCMV/LASgpc^2m^) and in guinea pigs challenged with guinea pig-adapted LASV (*26, 27*). Protection was achieved despite undetectable or low levels of neutralizing antibodies (nAbs), indicating that nAbs were not required for protection.

LASV has a bi-segmented single-stranded RNA genome. The small (S) segment encodes the nucleoprotein (NP) and GPC, whereas the large (L) segment encodes the viral polymerase (L) and matrix protein (Z). NP encapsidates viral genomic RNA to form the nucleocapsid, while GPC is processed into the stable signal peptide, GP1 and GP2, which form the trimeric GP complex responsible for receptor recognition and membrane fusion. Although the GP complex is the sole viral surface protein and the primary target of nAbs, nAb responses to LASV are generally weak and delayed in LASV-infected humans and animals (*15, 28*). Nonetheless, GPC has primarily been used as a vaccine antigen across LASV vaccine platforms, mainly because of its ability to elicit protective cellular immunity or non-nAbs depending on vaccine platform (*18*). The substantial genetic diversity of LASV represents a major challenge for the development of broadly protective vaccines (*29–31*). Notably, studies of LF survivors have demonstrated cross-lineage reactivity of LASV-specific T cells and identified conserved immunodominant T-cell epitopes in LASV GPC and NP (*32–34*), suggesting that the inclusion of NP may help broaden vaccine-induced cellular immunity beyond responses against GPC alone (*35, 36*). Experimental LASV vaccines expressing LASV NP have demonstrated varying degrees of success in preclinical models. Delivery via vaccinia virus (*37*) or Modified Vaccinia Ankara (MVA) (*38*) conferred full protection against fatal LF in guinea pigs, an effect seemingly mediated by robust, polyfunctional LASV-specific T-cell responses (*39*). However, the efficacy of the vaccinia-vectored vaccine expressing LASV NP was notably limited in rhesus macaques (*40*). Mucosal immunization with Salmonella typhimurium expressing LASV NP induced cross-protection against fatal LCMV challenge in mice (*41*), and LASV NP DNA-based immunization similarly protected mice against lethal LCMV challenge (*42*). Notwithstanding these findings, the protective potential of LASV NP antigen within mRNA vaccine formulations remains unknown and warrants further investigation. To overcome BSL-4 containment limitations and investigate the immunogenic potential of LASV NP and GPC in the context of the mRNA vaccine platform, we adapted a previously described lethal CBA/J-ML29 challenge model offering a suitable evaluation system outside a BSL-4 facility (*43*). In this model, the ML29 vaccine candidate, which carries LASV (lineage IV) S RNA and non-pathogenic Mopeia virus L RNA (*44, 45*), induces robust, LASV-specific protective T-cell immunity after parenteral immunization. However, intracerebral (i.c.) inoculations in mice induce a fatal, LCMV-like T-cell-mediated disease. This specific phenomenon has established a reliable small animal model for the efficacy evaluation of novel LASV vaccine candidates (*43, 46*).

Here, we evaluated two monovalent vaccines, an mRNA-LNP vaccine encoding LASV NP (LASnp-mRNA-LNP) and LASgpc-mRNA-LNP, and a bivalent vaccine combining the two. We characterized antigen-specific T-cell responses to the monovalent vaccines in CBA/N mice and evaluated the protective efficacy of both monovalent and bivalent vaccination in a lethal i.c. rML29 challenge model (CBA/N-rML29 model). We further evaluated the bivalent vaccine against lethal authentic LASV challenge in guinea pigs. The monovalent vaccines induced robust antigen-specific T-cell responses and, together with the bivalent formulation, conferred strong protection in the CBA/N-rML29 model. Importantly, low-dose bivalent vaccination provided complete protection against lethal LASV challenge in guinea pigs.

## RESULTS

### Monovalent LASgpc-mRNA-LNP and LASnp-mRNA-LNP confer protection in the CBA/N-rML29 model

*In vitro* transcribed (IVT) mRNAs encoding the LASV GPC or NP from the Josiah strain (lineage IV), with a 5′-cap and 3′-poly(A) tail, were generated using linearized DNA templates and 5-methoxyuridine-5′-triphosphate (5-methoxy-UTP) instead of UTP (Fig. S1). IVT mRNAs were then separately encapsulated in LNPs. Consistent with previous research using biologically derived ML29 in CBA/J mice (*43*), all CBA/N mice i.c. inoculated with ML29 generated via reverse genetics (rML29) (Fig. 1A) succumbed to infection within 10 days post-inoculation (Fig. S2). Using this model, mice were intramuscularly (i.m.) immunized with LASnp-mRNA-LNP or LASgpc-mRNA-LNP, each containing 2 µg of IVT mRNA, or mock-immunized twice at a 21-day interval and subsequently challenged i.c. with rML29 28 days later (Fig. 1B). All mock-immunized mice succumbed to infection within 8 days post-viral exposure (dpe), whereas all mice immunized with LASgpc-mRNA-LNP and four of five mice immunized with LASnp-mRNA-LNP survived (Fig. 1C). Antigen-specific antibody production was assessed by ELISA using plasma collected 14 days after the second immunization (Fig. S3). As previously reported (*26*), LASV GPC-specific antibodies were not detected (Fig. 1D). Likewise, NP-specific antibody responses were negligible (Fig. 1E). Because most LASV nAbs recognize conformational epitopes that require an assembled GP complex (*47*), neutralizing activity was examined by a focus reduction neutralizing test (FRNT). No neutralization activity was observed in plasma from mock- or LASgpc-mRNA-LNP-immunized mice (Fig. 1F). These results suggest that LASnp-mRNA-LNP and LASgpc-mRNA-LNP confer strong protection against lethal rML29 challenge in CBA/N mice despite the absence of detectable binding antibodies and nAbs.

**Fig. 1.**
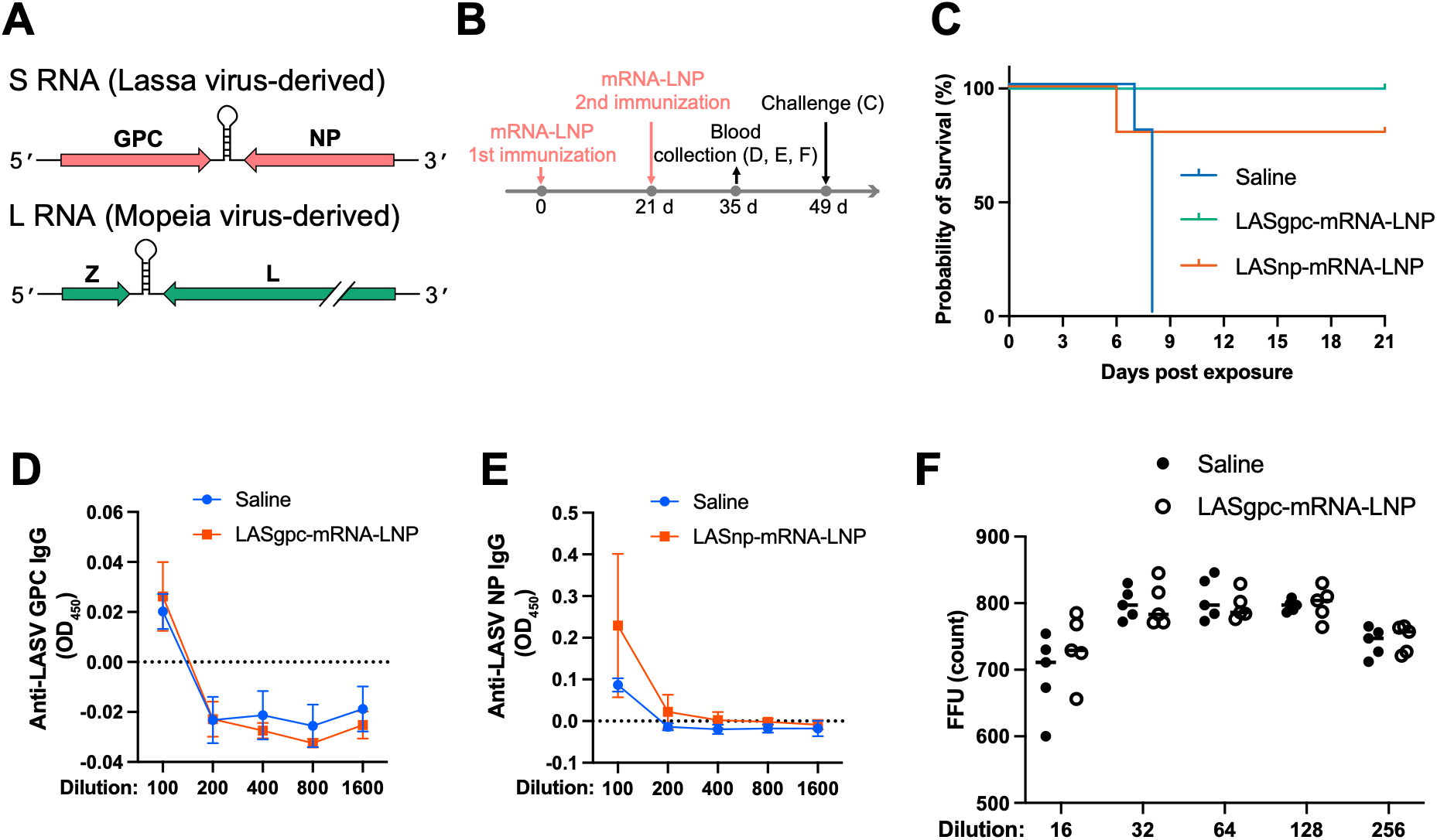
Efficacy evaluation of monovalent LASgpc-mRNA-LNP and LASnp-mRNA-LNP in the CBA/N mouse model of lethal rML29 infection. (**A**) Schematic representation of the ML29 genome. (**B**) Schematic overview of the experimental design for (**C**–**F**). CBA/N mice (n = 5 per group) were immunized twice with LASgpc-mRNA-LNP or LASnp-mRNA-LNP or mock-immunized with saline, 21 days apart. Fourteen days after the second immunization, blood samples were collected for antibody tests. Twenty-eight days after the second immunization, mice were inoculated i.c. with rML29. (**C**) Survival of CBA/N mice immunized with monovalent vaccine or mock-immunized following challenge with rML29. (**D**, **E**) Detection of LASV GPC-specific (**D**) and NP-specific (**E**) antibodies in plasma from LASgpc-mRNA-LNP-, LASnp-mRNA-LNP-, or mock-immunized mice by ELISA. OD values from samples without plasma or antibodies were subtracted as background. (**F**) Neutralizing activity of plasma from LASgpc-mRNA-LNP-immunized or mock-immunized mice determined by FRNT.

### Fine mapping of LASV GPC-specific CD4^+^ and CD8^+^ T-cell epitopes in CBA/N mice

LASgpc-mRNA-LNP protected CBA/N mice against lethal rML29 exposure without inducing detectable nAb responses, suggesting that protection was mediated by GPC-specific T-cell responses. Six LASV GPC-derived 20-mer peptides spanning amino acid (aa) positions 57–83, 218–244, and 400–426 were previously shown to be recognized by T cells from ML29-immunized CBA/J mice (*43*). To better characterize vaccine-induced T-cell responses, we performed fine mapping of GPC-specific CD4^+^ and CD8^+^ T-cell epitopes. CBA/N mice were intraperitoneally (i.p.) inoculated with rML29, and splenocytes were isolated 8 days later. To identify CD8^+^ T-cell epitopes, 57 9-mer peptides with a 1-aa offset derived from these peptides were individually used to stimulate splenocytes. IFN-γ production was assessed by ELISPOT assay, and peptides GPC-9 and GPC-10 yielded markedly high spot counts (Fig. 2A). Testing peptides of different lengths (8– 10 aa) within the region covered by GPC-9 and GPC-10 (aa 65–74) showed that the 8-mer peptide and both 9-mer peptides, but not the 10-mer peptide, induced robust IFN-γ production (Fig. 2B). The 8-mer peptide GPC_66-73_, the shortest reactive peptide tested, was selected for subsequent experiments. Despite the strong IFN-γ production reportedly induced by two 20-mer peptides spanning aa 218–244 (GPC_218-237_ and GPC_225-244_) in a previous study (*43*), no 9-mer peptides covering the same region yielded high spot counts (Fig. 2A), suggesting that this region contains a CD4^+^ T-cell epitope that may not be efficiently presented by 9-mer peptides. To address this, we tested two 20-mer peptides and three 15-mer peptides (GPC_223-237,_ GPC_224-238_, and GPC_225-239_) with a 1-aa offset containing the 13-aa sequence shared by GPC_218-237_ and GPC_225-244_. These peptides were used to stimulate splenocytes from rML29-immune CBA/N mice, with or without the depletion of CD4⁺ or CD8⁺ cells. The two 20-mer peptides and all three 15-mer peptides yielded high spot counts, with GPC_225-239_ showing the highest mean spot counts (Fig. 2C). IFN-γ responses were abolished by CD4^+^ cell depletion but not by CD8^+^ cell depletion, indicating that these responses were CD4⁺ T cell-dependent (Fig. 2C). To validate the ability of GPC_225-239_ and GPC_66-73_ to stimulate IFN-γ production by LASV GPC-specific T cells, these peptides were used to stimulate splenocytes from mock- and rML29-immunized CBA/N mice, followed by intracellular cytokine staining (ICS). Stimulation with GPC_225-239_ and GPC_66-73_ significantly increased the frequencies of IFN-γ-producing CD4^+^ and CD8^+^ T cells, respectively, in the rML29-immunized group (Fig. 2D).

**Fig. 2.**
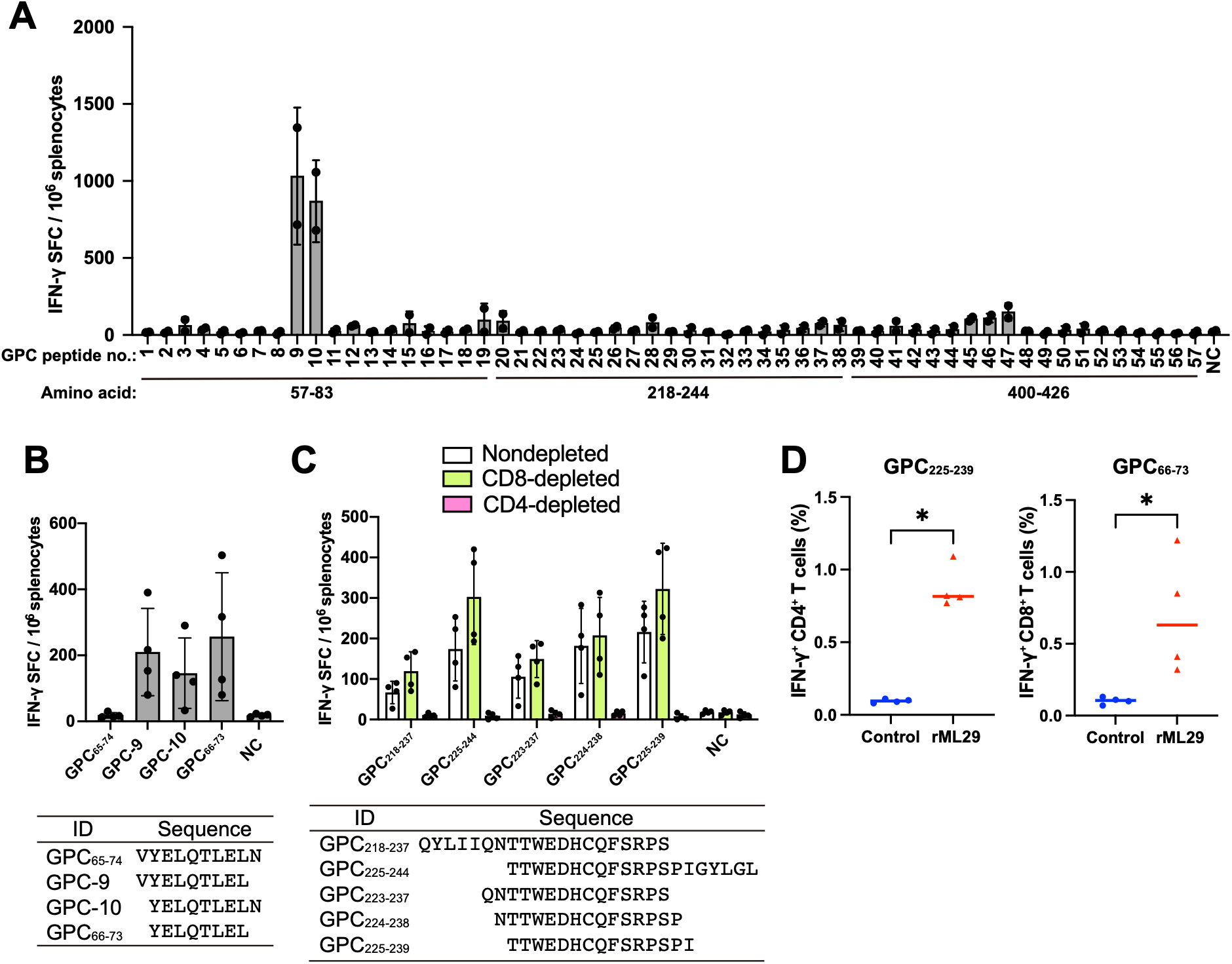
Identification of immunodominant T-cell epitopes of LASgpc. (**A**) IFN-γ-producing cells (spot-forming cells, SFCs) were quantified by ELISPOT assay after stimulation of splenocytes from rML29-immunized CBA/N mice with LASV GPC-derived 9-mer peptides. (**B**) IFN-γ secretion by splenocytes from rML29-immunized CBA/N mice after stimulation with selected 8–10-mer peptides spanning the region covered by the peptides GPC-9 and GPC-10 determined by ELISPOT. The peptide sequences are listed below. (**C**) Quantification of IFN-γ SFCs by ELISPOT assay after stimulation of nondepleted, CD8⁺-depleted, or CD4⁺-depleted splenocytes from rML29-immunized CBA/N mice with 15- or 20-mer peptides with the indicated sequences. (**D**) Frequencies of IFN-γ-producing cells within the CD4^+^ or CD8^+^ T-cell populations in splenocytes from rML29-immunized CBA/N mice following stimulation with GPC_225–239_ or GPC_66–73_ determined by flow cytometry. Statistical significance: *P < 0.05, Mann–Whitney U test. Each dot represents an individual mouse. Data are shown as means ± SD in (**A**–**C**), and horizontal bars indicate means in (**D**).

### Identification of LASV NP-specific CD4^+^ and CD8^+^ T-cell epitopes in CBA/N mice

To investigate T-cell responses associated with LASnp-mRNA-LNP-mediated protection, we identified LASV NP-specific T-cell epitopes in CBA/N mice. We used an overlapping peptide (OLP) library composed of 15-mer peptides spanning the full-length LASV NP with a 4-aa offset. The OLPs were arranged into 24 peptide pools (Pools A–X) by a matrix pooling strategy, with each pool containing 8–12 peptides, and tested by ELISPOT assay using splenocytes from rML29-immunized CBA/N mice (Fig. 3A). Pools D, I, Q, and U yielded markedly higher IFN-γ-positive spot counts than the other peptide pools (Fig. 3B). Four peptides identified at the intersections of these pools (NP-41, NP-45, NP-101, and NP-105) were analyzed by ELISPOT with or without the depletion of CD4⁺ or CD8⁺ cells. NP-41 and NP-105, but not NP-45 or NP-101, showed high spot counts in nondepleted splenocytes, with higher mean counts for NP-41 than NP-105 (Fig. 3C). CD4^+^ cell depletion abolished the IFN-γ responses to NP-41 and NP-105, whereas CD8^+^ cell depletion did not, suggesting that these peptides contain CD4⁺ T-cell epitopes (Fig. 3C). These results also suggest that CD8⁺ T-cell epitopes may not be efficiently presented by these 15-mer OLPs. To identify CD8⁺ T-cell epitopes, we performed *in silico* analyses. H-2K^k^-restricted CD8^+^ T-cell epitopes were predicted using 10 different algorithms, and the top 3 candidates from each algorithm were extracted, resulting in the selection of 16 peptides (NP-CD8-1 to NP-CD8-16) for subsequent tests (Table 1). ELISPOT assays using these candidate peptides, except for NP-CD8-4, which could not be dissolved in DMSO, revealed that stimulation with NP-CD8-1 yielded markedly higher IFN-γ^+^ spot counts than the other peptides (Fig. 3D). ICS confirmed significant increases in the frequencies of IFN-γ^+^ CD4^+^ and IFN-γ^+^ CD8^+^ T cells in splenocytes from rML29-immunized CBA/N mice following stimulation with NP-41 and NP-CD8-1, respectively (Fig. 3E). Hereafter, NP-41 and NP-CD8-1 are referred to as NP_161-175_ and NP_303-310_, respectively, corresponding to their aa positions.

**Fig. 3.**
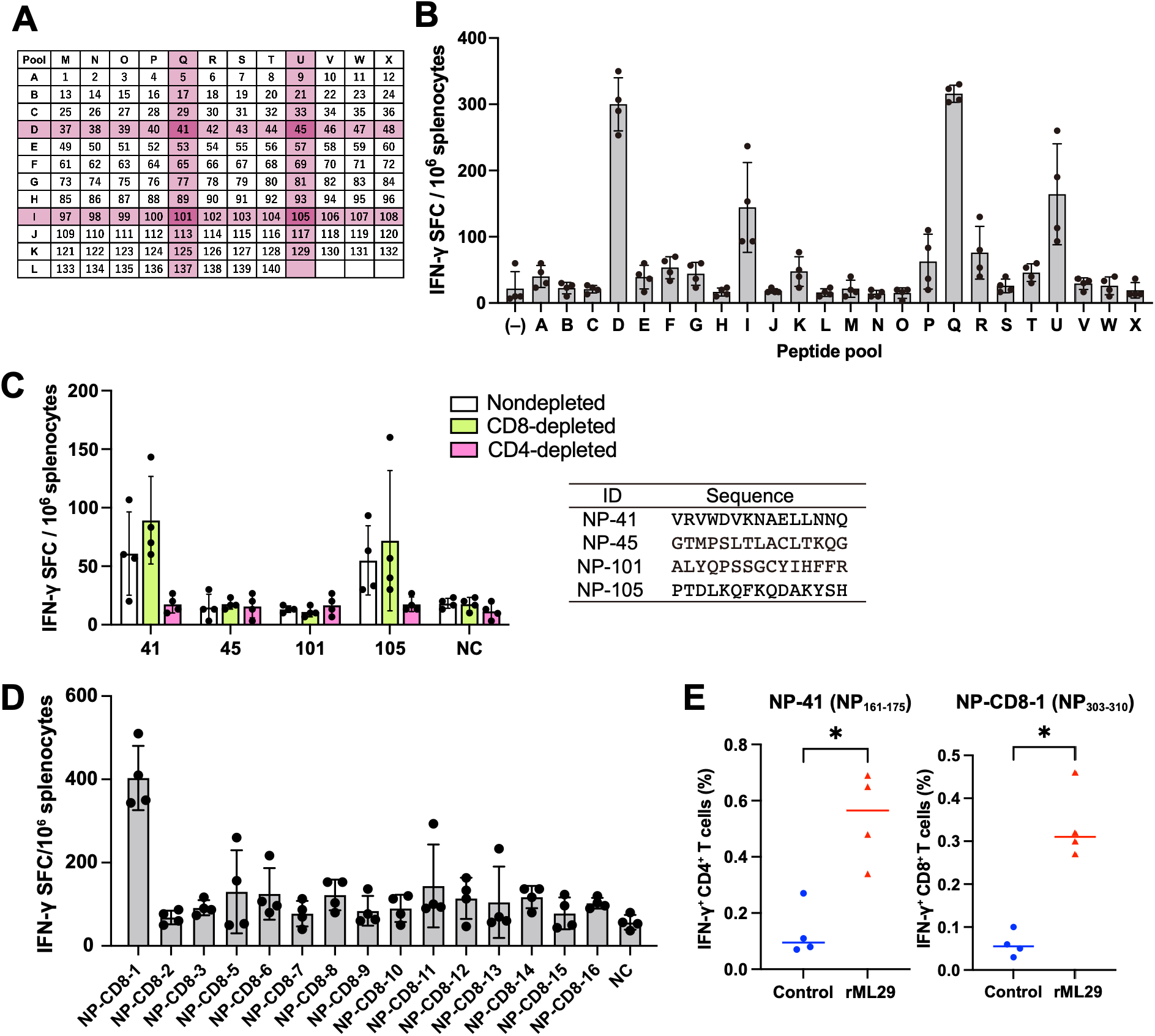
Identification of immunodominant T-cell epitopes of LASnp. (**A**) Arrangement of 24 peptide pools (A–X) generated from an overlapping peptide library composed of 140 15-mer peptides derived from LASV NP (NP-1–NP-140). Responsive peptide pools identified in (**B**) are highlighted in pink. (**B**) IFN-γ-producing cells (SFCs) were quantified by ELISPOT assay after stimulation of splenocytes from rML29-immunized CBA/N mice with the peptide pools (A–X). (**C**) Quantification of IFN-γ SFCs by ELISPOT assay after stimulation of nondepleted, CD8⁺-depleted, or CD4⁺-depleted splenocytes from rML29-immunized CBA/N mice with peptides located at the intersections of responsive pools identified in (**B**). The peptide sequences are listed. (**D**) Quantification of IFN-γ SFCs by ELISPOT assay after stimulation of splenocytes from rML29-immunized CBA/N mice with candidate LASV NP-specific CD8^+^ T-cell epitope peptides selected by *in silico* analysis (Table 1). (**E**) Frequencies of IFN-γ-producing cells within the CD4^+^ or CD8^+^ T-cell populations in splenocytes from rML29-immunized CBA/N mice following stimulation with NP-41 (NP_161-175_) or NP-CD8-1 (NP_303-310_) determined by flow cytometry. Statistical significance: *P < 0.05, Mann–Whitney U test. Each dot represents an individual mouse. Data are shown as means ± SD (**B**–**D**), and horizontal bars indicate means in (**E**).

**Table 1.**
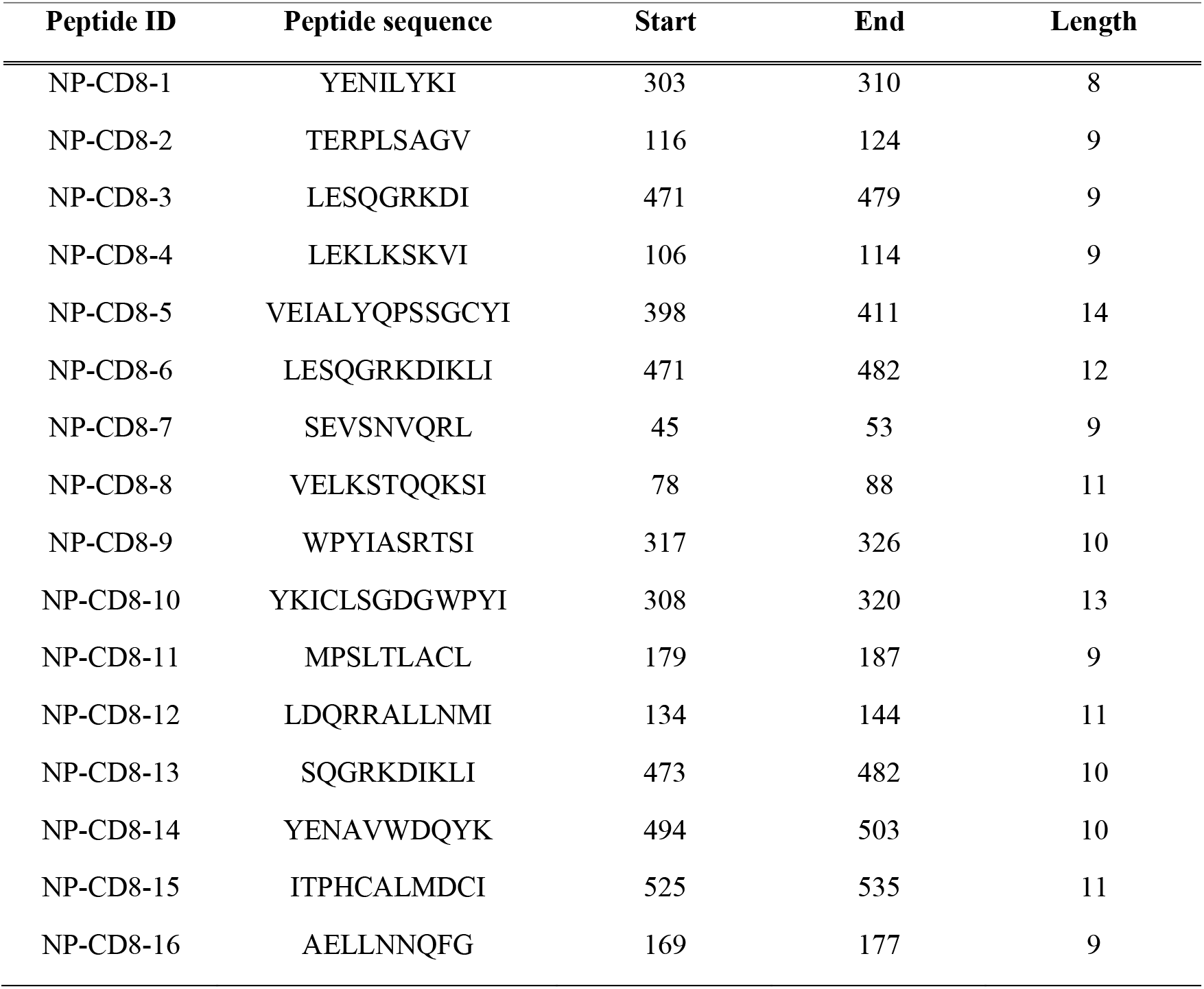
Candidate H-2K^k^-restricted LASnp-specific CD8⁺ T-cell epitopes predicted using the Immune Epitope Database (IEDB) T Cell Epitope Prediction tool.

### LASgpc-mRNA-LNP and LASnp-mRNA-LNP induce antigen-specific T-cell responses

Based on the described epitope mapping, we evaluated antigen-specific T-cell responses elicited by the monovalent mRNA vaccines. CBA/N mice were immunized i.m. with LASgpc-mRNA-LNP or LASnp-mRNA-LNP, each containing 2 µg of IVT mRNA, or mock-immunized twice at a 21-day interval. Eight days after the second immunization, splenocytes were isolated and stimulated with the epitope peptides GPC_225-239_, GPC_66-73_, NP_161-175_, and NP_303-310_, followed by ICS, to detect IFN-γ production (Fig. 4A). Immunization with LASgpc-mRNA-LNP significantly increased the frequencies of IFN-γ^+^ CD4^+^ and IFN-γ^+^ CD8^+^ T cells compared with those in the mock-immunized group (Fig. 4B). Mice immunized with LASnp-mRNA-LNP showed a significant increase in the frequency of IFN-γ^+^ CD8^+^ T cells, while IFN-γ^+^ CD4^+^ T cells also increased significantly, albeit modestly (Fig. 4C). Collectively, the absence of detectable nAb production suggests that antigen-specific T-cell responses drive the protection conferred by LASgpc-mRNA-LNP and LASnp-mRNA-LNP against lethal rML29 infection. Furthermore, aa sequence alignment across various LASV lineages revealed a high degree of conservation for the GPC- and NP-specific CD4^+^ and CD8^+^ T-cell epitopes identified in CBA/N mice (Fig. 4D). These findings suggest that the epitope mapping performed in this study provides a valuable framework for the future evaluation of vaccines targeting LASV lineages beyond lineage IV within the CBA/N mouse model.

**Fig. 4.**
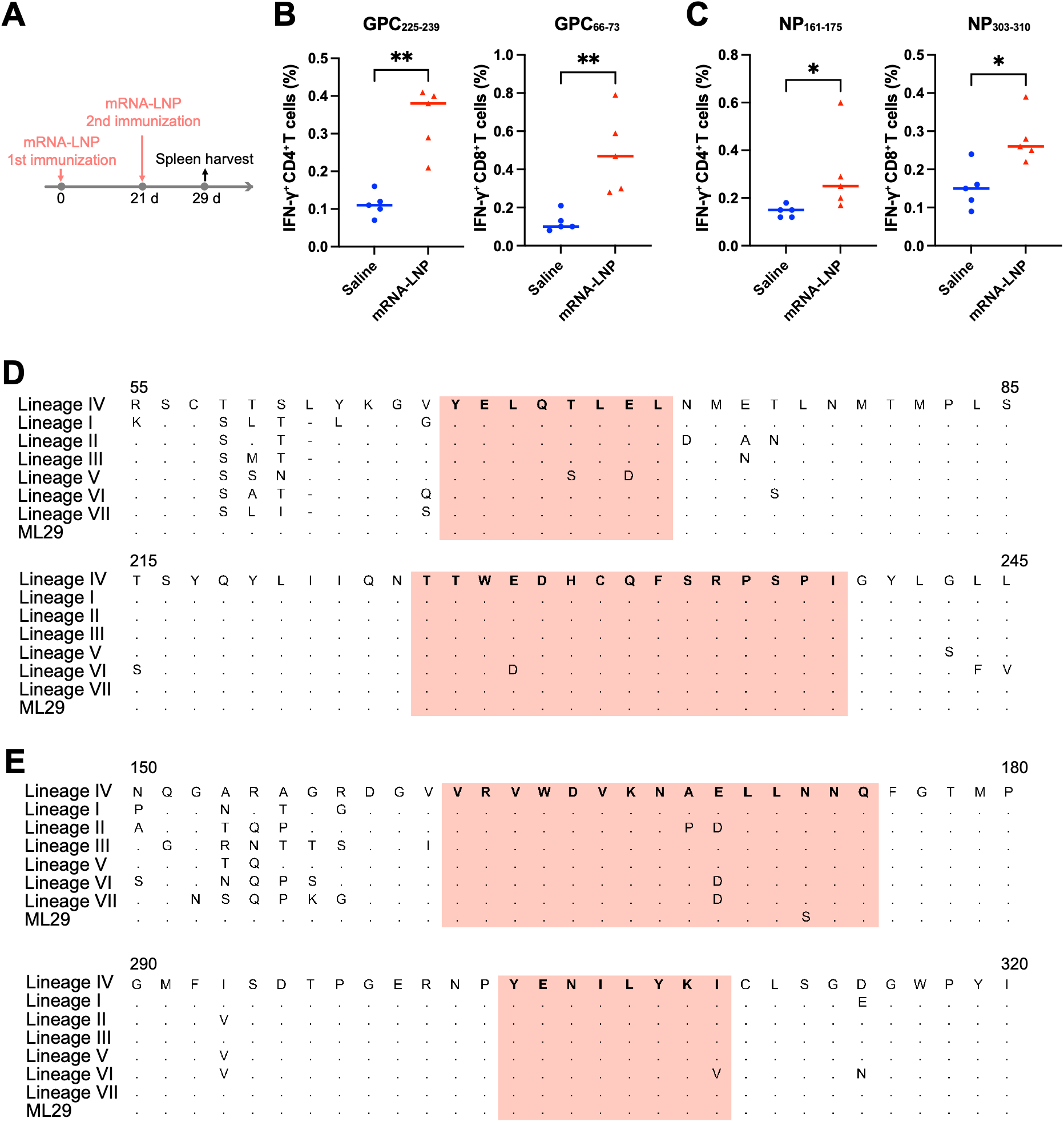
Antigen-specific T-cell responses in CBA/N mice immunized with LASgpc-mRNA-LNP or LASnp-mRNA-LNP. (**A**) Schematic overview of the experimental design for (**B** and **C**). CBA/N mice were immunized twice with LASgpc-mRNA-LNP or LASnp-mRNA-LNP or mock-immunized with saline, 21 days apart. Eight days after the second immunization, spleens were harvested to assess antigen-specific T-cells responses (**B**, **C**) Frequencies of IFN-γ-producing cells within the CD4^+^ or CD8^+^ T-cell populations in splenocytes from CBA/N mice twice immunized with a monovalent vaccine or mock-immunized with saline following stimulation with the indicated peptides determined by flow cytometry. Statistical significance: **P < 0.01, *P < 0.05, Mann–Whitney U test. Each dot represents an individual mouse. Horizontal bars indicate means. (**D**, **E**) Amino acid sequence alignments across LASV lineages of regions containing LASV GPC-specific (**D**) and NP-specific (**E**) T-cell epitopes identified in CBA/N mice. The lineage IV sequence was used as the reference. Amino acids identical to the reference sequence are indicated by dots. The regions corresponding to the CD4^+^ and CD8^+^ T-cell epitope peptides identified in this study are highlighted in salmon pink. LASV GPC sequence information was obtained from NCBI: lineage IV, Josiah, NP_694870.1; lineage I, Pinneo1969, AIT17836.1; lineage II, Nig08-04, ADU56610.1; lineage III, Nig08-A19, ADU56618.1; lineage V, Soromba-R, AHC95553.1; lineage VI, KAK-428, ANH09740.1; lineage VII, Alzey, SCA79105.1; ML29, YP_170705.1. LASV NP sequence information was obtained from NCBI: lineage IV, Josiah, NP_694869.1; lineage I, Pinneo1969, AIT17837.1; lineage II, Nig08-04, ADU56611.1; lineage III, Nig08-A19, ADU56619.1; lineage V, Soromba-R, AHC95552.1; lineage VI, KAK-428, ANH09741.1; lineage VII, Alzey, SCA79106.1; ML29, YP_170706.1.

### Bivalent mRNA-LNP vaccine has enhanced protective efficacy in the CBA/N-rML29 model

LASgpc-mRNA-LNP and LASnp-mRNA-LNP provided high levels of protection in the CBA/N-rML29 model (Fig. 1C). We therefore investigated the protective efficacy of a bivalent formulation combining the two monovalent mRNA vaccines. CBA/N mice were mock-immunized or immunized twice at a 21-day interval with monovalent LASgpc-mRNA-LNP or LASnp-mRNA-LNP, each containing 0.4 µg of IVT mRNA, or with a bivalent vaccine combining LASgpc-mRNA-LNP and LASnp-mRNA-LNP, each containing 0.2 µg of IVT mRNA. The total mRNA dose was 0.4 µg in the monovalent and bivalent vaccine groups. Twenty-eight days after the second immunization, mice were challenged i.c. with rML29 to assess protective efficacy (Fig. 5A). At this reduced dose, the protective efficacy of monovalent vaccines was lower than that observed at a 2-µg dose (Fig. 1C). Notably, partial protection was observed in the monovalent cohorts, where one of five mice immunized with LASgpc-mRNA-LNP and four of five mice immunized with LASnp-mRNA-LNP succumbed to rML29 infection (Fig. 5B). In striking contrast, the bivalent formulation conferred complete protection against lethal rML29 challenge across all immunized mice, despite utilizing only half the individual dose (0.2 µg) of each constituent mRNA compared to the monovalent formulations. These findings suggest that the bivalent vaccine platform may provide enhanced protective efficacy compared to monovalent approaches, potentially through complementary immunological mechanisms.

**Fig. 5.**
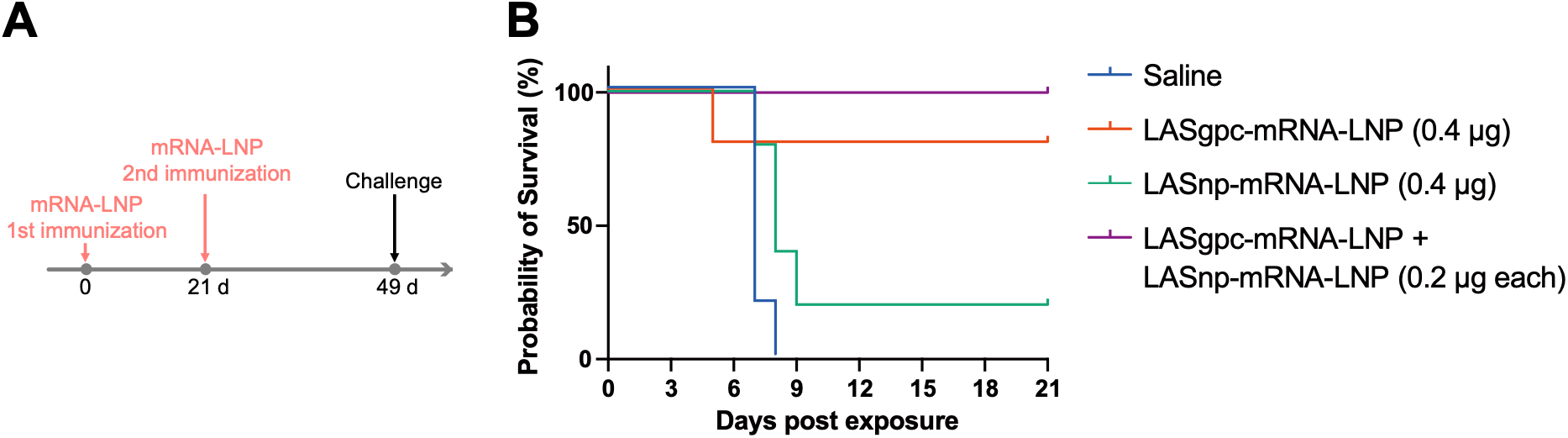
Efficacy evaluation of a bivalent vaccine combining LASgpc-mRNA-LNP and LASnp-mRNA-LNP in the CBA/N-rML29 challenge model. (**A**) Schematic overview of the experimental design for (**B**). CBA/N mice (n = 5 per group) were immunized twice with LASgpc-mRNA-LNP (0.4 µg mRNA), LASnp-mRNA-LNP (0.4 µg mRNA), a bivalent vaccine combining the two (0.2 µg mRNA each), or mock-immunized with saline, 21 days apart. Twenty-eight days after the second immunization, mice were inoculated i.c. with rML29. (**B**) Survival of CBA/N mice following lethal challenge with rML29 after immunization with the indicated monovalent or bivalent mRNA-LNP vaccines.

### Bivalent vaccination fully protects guinea pigs from lethal LASV challenge

Finally, we investigated whether the high levels of protection conferred by the bivalent vaccine in the CBA/N-rML29 model could be extended to a lethal guinea pig model of authentic LASV infection (*48*) using refined vaccine preparations. For this, IVT mRNAs encoding codon-optimized LASV GPC and NP were encapsulated into SM102-based LNPs, generating optLASgpc-mRNA-LNP and optLASnp-mRNA-LNP, respectively. In preliminary experiments, protein expression from the newly synthesized IVT mRNAs and protective efficacy of a bivalent formulation combining the two at high (1 µg mRNA each) and low (0.2 µg mRNA each) doses in the CBA/N-rML29 model were confirmed in Western blot and challenge experiments, respectively (Fig. S4). Guinea pigs were mock-immunized or immunized, twice at a 21-day interval, with the bivalent vaccine combining optLASgpc-mRNA-LNP and optLASnp-mRNA-LNP at a high (5 µg mRNA each for the first immunization and 2.5 µg mRNA each for the second immunization) or low (1 µg mRNA each) dose. No overt adverse effects were observed after vaccination (Fig. S5). Twenty-eight days after the second immunization, guinea pigs were challenged with a lethal dose of a non-guinea-pig-adapted clinical isolate of LASV (lineage IV, strain LF2384) (Fig. 6A). Four of five mock-immunized guinea pigs succumbed to LASV infection. In contrast, all guinea pigs immunized with the high or low dose of the bivalent formulation survived (Fig. 6B). Mock-immunized guinea pigs showed gradual weight loss from 10 dpe, and transiently elevated body temperatures (>40°C) from 9 to 16 dpe, whereas the high-and low-dose groups showed no abnormal changes in body weight or temperature (Fig. 6C). To determine whether antigen-specific antibody responses were induced by the bivalent vaccine, sera collected 14 days after the second immunization were analyzed by ELISA. In contrast to the results obtained in CBA/N mice (Fig. 1D, E), antigen-specific antibodies against LASV GPC and NP were readily detected in guinea pigs vaccinated with the bivalent vaccine at high and low doses (Fig. 6D). Next, we evaluated whether these GPC-specific antibody responses were associated with neutralizing activity. Consistent with our previous observations (Fig. 1F), no neutralizing activity was detected in the sera from bivalent vaccine-immunized or mock-immunized animals (Fig. 6E). These results demonstrate that the bivalent mRNA-LNP vaccine conferred robust protection against lethal LASV infection in guinea pigs, even in the absence of detectable nAb responses.

**Fig. 6.**
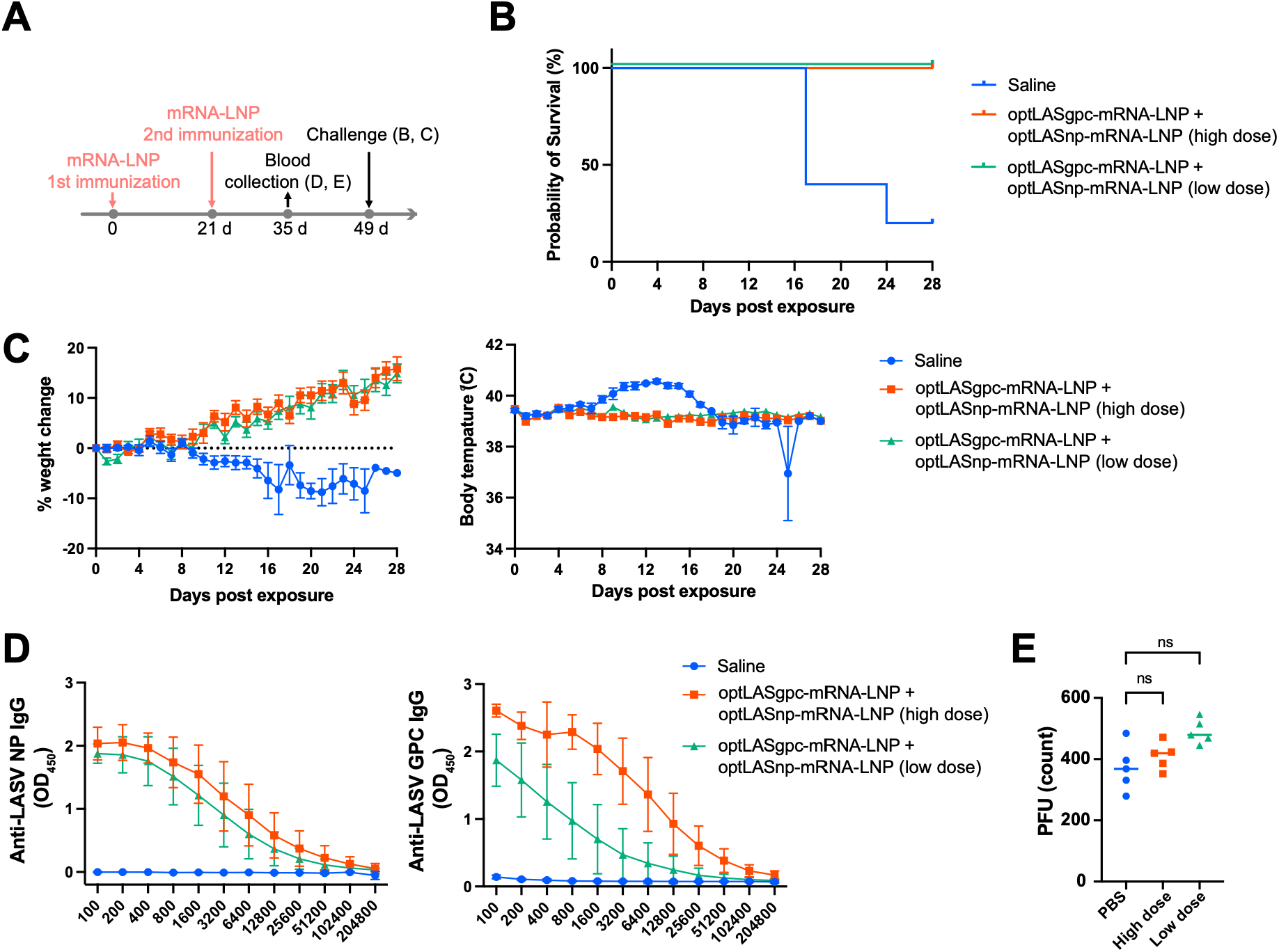
Efficacy evaluation of a bivalent vaccine combining optLASgpc-mRNA-LNP and optLASnp-mRNA-LNP in a lethal guinea pig model of LASV infection. (A) Schematic overview of the experimental design for (B–E). Hartley guinea pigs (n = 5 per group) were immunized twice with a bivalent vaccine combining optLASgpc-mRNA-LNP and optLASnp-mRNA-LNP at a high (5 µg mRNA each for the first immunization and 2.5 µg mRNA each for the second immunization) or low (1 µg mRNA each) dose or mock-immunized with saline, 21 days apart. Fourteen days after the second immunization, blood samples were collected for antibody tests. Twenty-eight days after the second immunization, guinea pigs were inoculated i.p. with LASV. (B, C) Survival (B), body weight (C, left), and body temperature (C, right) of guinea pigs following LASV challenge. (**D**) Detection of LASV GPC-specific (left) and NP-specific (right) antibodies in serum from guinea pigs immunized with the bivalent vaccine at high or low doses or mock-immunized with saline by ELISA. (**E**) Neutralizing activity of serum from the same groups, as determined by PRNT. Statistical significance: ns; nonsignificant, Mann–Whitney U test. Data are shown as means ± SD (**C**, **D**). In (**E**), each dot represents an individual guinea pig and horizontal bars indicate means.

## DISCUSSION

We previously demonstrated the protective efficacy of LASgpc-mRNA-LNPs in a lethal rLCMV/LASgpc^2m^ infection model in CBA/N mice (*26*). Ronk A. J. et al. also reported that LASV GPC mRNA-LNP vaccine protected guinea pigs against fatal LF independently of nAb responses (*27*). Here, we confirmed this observation in the CBA/N-rML29 model and documented for the first time partial protective efficacy of LASnp-mRNA-LNP. NP is the most abundant viral protein produced in infected cells, rendering it a primary target for both T-cell-mediated immunity and non-nAbs. Prior studies have demonstrated that cytotoxic T-cell responses directed against early NP-derived epitopes are significantly more efficacious in viral control than those targeting late GPC-derived epitopes (*49*). Historically, the first experimental LASV vaccine, a vaccinia virus vector expressing LASV NP (*37, 40*), conferred partial protection in rhesus macaques and complete protection in guinea pig models, underscoring the clinical relevance of NP-based vaccine strategies. In this study, we demonstrate that immunization with LASnp- and LASgpc-mRNA-LNP experimental vaccines confers robust protection in both murine model of rML29 infection and guinea pig model of LASV infection. Notably, the bivalent formulation achieved complete protection against rML29 lethal challenge in CBA/N mice, even at mRNA-LNP dosage where monovalent counterparts failed to protect all mice, and against a lethal challenge with the clinical LASV isolate LF2384 in guinea pigs. These findings confirm the efficacy of both LASV GPC and NP as critical antigens for LASV mRNA vaccine development and establish a compelling rationale for the development of bivalent mRNA-LNP strategies to achieve dose sparing while maintaining protection against LF.

The CBA/J mouse model of i.c. ML29 challenge system has been successfully applied to assess experimental LASV vaccine candidates, including ML29, alphavirus RNA replicon-based constructs, and MVA-vectored vaccines, in a cost-effective manner outside BSL-4 containment (*43, 46*). Because protection in this model is mediated primarily by virus-specific T-cell responses, we performed fine mapping of LASV GPF and NP-specific CD4^+^ and CD8^+^ T-cell epitopes in CBA/N mice. Intriguingly, rML29 immunization in CBA/N mice induced comparable CD4⁺ and CD8⁺ T-cell responses, whereas LASgpc-mRNA-LNP and LASnp-mRNA-LNP predominantly elicited CD8⁺ T-cell responses. Furthermore, LASV NP-specific CD4⁺ T-cell responses were particularly limited following LASnp-mRNA-LNP immunization. A better understanding of the mechanisms underlying the limited CD4^+^ T-cell responses elicited by LASnp-mRNA-LNP may help inform strategies to achieve the balanced T-cell immunity observed following rML29 immunization, which may improve vaccine efficacy and durability, potentially reducing the number of immunizations required for protection against LF.

Neither LASgpc-mRNA-LNP nor LASnp-mRNA-LNP induced detectable antigen-specific antibodies in CBA/N mice, whereas robust responses against LASV GPC and NP were observed in guinea pigs immunized with the bivalent vaccine. However, neutralizing activity was not detected in either model. These observations on GPC-specific antibody responses are consistent with previous observations in ML29-immunized CBA/J mice (*43*) and in guinea pigs in a LASV mRNA vaccine study (*27*). The original ML29 challenge/vaccination model was established using the CBA/J substrain (43). In the present study, we adapted this system by employing rML29 rescued from cDNA clones together with the CBA/N substrain. Although CBA/N mice retain the capacity to produce antigen-specific antibodies to certain antigens (*50*), they carry the X-linked immunodeficiency (xid) mutation and exhibit impaired B-cell responses depending on the antigen and immunization conditions (*51*). Thus, we cannot exclude the possibility that the xid-associated B-cell defect contributed to the lack of detectable GPC- and NP-specific antibody responses observed in CBA/N mice. In addition, the discrepancy in antigen-specific antibody production between mice and guinea pigs may reflect differences in host species, mRNA-LNP formulation, immunization dose, antigen expression, ELISA antigen, and assay sensitivity. Thus, the lack of detectable LASV antigen-specific antibodies in mice does not necessarily indicate the intrinsic inability of LASV mRNA-LNP vaccines to induce humoral responses. These findings underscore the importance of evaluating humoral and cellular immune responses in multiple animal models when evaluating LASV vaccine candidates. Nonetheless, the CBA/N-rML29 model provides a suitable experimental system for LASV vaccine research because these mice can develop robust LASV antigen-specific T cell responses, which can be rapidly characterized in detail using the T-cell epitope information identified in this study, and because ML29 infection can be performed outside a BSL-4 facility. Therefore, our two-step approach—initial assessment and optimization of LASV vaccine candidates using the CBA/N-rML29 model, followed by validation of selected candidates in the lethal guinea pig model of LASV infection—can substantially mitigate the challenges associated with the early stages of LASV vaccine research, including the need for a BSL-4 facility and the use of guinea pigs, for which an evaluation system of T-cell responses has not been well established.

Owing to the extensive genetic diversity among LASV lineages, the ability of a single vaccine to confer cross-lineage protection is an important consideration in LASV vaccine development. LASV GPC- and NP-specific T-cell responses have been reported to exhibit broad cross-reactivity across genetically diverse LASV strains (*32, 33*). Indeed, several LAV candidates expressing both antigens induced cross-reactive protective immunity in preclinical studies (*36, 52*). Intriguingly, a measles virus-vectored vaccine candidate expressing LASV GPC and NP exhibited greater efficacy than one expressing GPC and Z (*35*). Therefore, including both antigens represents an important strategy for conferring protection against a broader range of lineages, as well as for enhancing efficacy. Consistent with these observations, we found that the co-administration of LASgpc-mRNA-LNP and LASnp-mRNA-LNP provided complete protection at lower doses than either monovalent vaccine in the CBA/N-rML29 model and also conferred complete protection against lethal LASV challenge in guinea pigs at 1 µg mRNA per antigen, a dose substantially lower than the dose reported previously (*27*), although differences in construct, formulation, and challenge virus preclude direct comparison. Notably, in rhesus macaques immunized with a vaccinia virus construct simultaneously expressing all S segment-encoded LASV proteins (V-LSG/N), LASV was only barely detectable in tissues after a fatal challenge, in contrast to animals that received individually expressed antigens (V-LSG or V-LSN) (*40*).

This study had several limitations. First, immune correlates of protection were not determined. Although our data demonstrated that LASgpc-mRNA-LNP and LASnp-mRNA-LNP elicited robust antigen-specific T-cell responses in mice prior to rML29 challenge, we did not directly evaluate the contribution of CD4⁺ or CD8⁺ T cells to vaccine-mediated protection. In addition, T-cell responses in guinea pigs were not evaluated because of the limited availability of immunological assays for this species. Future studies involving the adoptive transfer of T cells from vaccinated animals are needed to define the relative contribution of each T-cell subset to vaccine-mediated protection, particularly in the context of bivalent vaccination. Second, the lack of *in vitro* neutralizing activity does not necessarily indicate that humoral responses did not contribute to protection. Non-nAbs may mediate antiviral protection through Fc-dependent effector functions, including antibody-dependent cellular cytotoxicity (ADCC) and antibody-dependent cellular phagocytosis (ADCP), as demonstrated with LASSARABV, an inactivated LASV vaccine candidate based on recombinant rabies virus expressing LASV GPC (*53*). Passive transfer experiments using serum from vaccinated animals, together with *in vitro* assessments of ADCC and ADCP activities, might help clarify the role of humoral immunity in protection. Finally, single-dose efficacy and durability are key attributes for LASV vaccine candidates as emphasized in the WHO TPP. Because these aspects are not evaluated in the present study, assessing protective efficacy following a single immunization and the durability of immune responses will be an important next step toward clinical development.

In this study, the bivalent LASV mRNA vaccine conferred strong protection in the guinea pig model of LASV infection at a dose of 1 µg mRNA per antigen. The surrogate CBA/N-rML29 model, together with the T-cell epitope information obtained in this study, provides a practical platform for vaccine optimization and mechanistic studies, thereby reducing the workload at BSL-4 facilities. Future studies should utilize this platform to optimize vaccine formulations, followed by evaluation of selected vaccine candidates for the durability and breadth of protection against homologous and genetically distant heterologous strains in authentic LASV infection models. These studies will provide an important foundation for advancing bivalent mRNA-LNP vaccines against LF toward clinical development.

## MATERIALS AND METHODS

### mRNA synthesis and LNP preparation used in the CBA/N-rML29 model

DNA fragments encoding LASV GPC or NP open reading frame (ORF) from lineage IV Josiah strain (NCBI accession no. NC_004296.1) were inserted into the Linearized Template Vector provided with the Takara IVTpro Synthesis System (Takara, Shiga, Japan). A synonymous mutation was introduced into the NP ORF to disrupt the *Sac*I restriction site. The resulting plasmids were digested with *Hin*dIII and *Sac*I, and template DNA fragments containing either the GPC or NP ORF were isolated by agarose gel electrophoresis and gel extraction. IVT mRNAs were synthesized using the Takara IVTpro Synthesis System (Takara) according to the manufacturer’s instructions with minor modifications. Briefly, 5-methoxy-UTP (TriLink BioTechnologies, San Diego, CA, USA) was used instead of UTP, together with CleanCap Reagent AG (TriLink Bio Technologies). Synthesized mRNAs were then purified using the Monarch RNA clean up kit (New England Biolabs, Ipswich, MA, USA), and the quality of the purified IVT mRNAs was assessed by non-denaturing agarose gel electrophoresis using RNA High for Easy Electrophoresis (DynaMarker Laboratory, Tokyo, Japan). LNPs were prepared using GenVoy-ILM kit (NanoAssemblr Ignite Training Kit with GenVoy-ILM, Precision Nanosystems, Vancouver, BC, Canada) according to the manufacturer’s recommendations. IVT-mRNA-LNPs were formulated by mixing GenVoy-LNP (Precision Nanosystems) and IVT mRNA using a microfluidic mixer (NanoAssemblr Ignite; Precision Nanosystems). The resulting samples were subsequently diluted with 1 × Formulation Buffer 2 (Precision Nanosystems), concentrated using Amicon Ultra centrifugal filters (10-kDa cutoff; Merck, Darmstadt, Germany), and filtered through a 0.2-µm filter (Pall, Port Washington, NY, USA). The sterile-filtered IVT-mRNA-LNPs were stored at 4°C until use. The concentration of IVT-mRNA incorporated into the LNPs was quantified using a Quant-iT RiboGreen RNA Assay Kit (Thermo Fisher Scientific, Waltham, MA, USA).

### mRNA synthesis and LNP preparation used in the guinea pig model of LASV infection

5-Methoxyuridine-modified mRNAs encoding codon-optimized LASV GPC (optLASgpc-mRNA) or NP (optLASnp-mRNA) were synthesized by ARCALIS (Kashiwa, Chiba, Japan). Heptadecan-9-yl 8-[(2-hydroxyethyl)(6-oxo-6-(undecyloxy)hexyl)amino]octanoate (SM-102) was purchased from SHINOPEG (Xiamen, China). 1,2-Distearoyl-sn-glycero-3-phosphocholine (DSPC), cholesterol, and 1,2-dimyristoyl-rac-glycero-3-methoxypolyethylene glycol-2000 (DMG-PEG2K) were purchased from NOF Corporation (Tokyo, Japan). mRNA-LNPs were prepared using SM-102, DSPC, cholesterol, and DMG-PEG2K at a molar ratio of 48.4:11.3:37.8:2.5, respectively. The lipids were dissolved in ethanol. optLASgpc-mRNA and optLASnp-mRNA were each prepared in 50 mM citrate buffer (pH 5.0). The lipid solution and mRNA solution were mixed using a microfluidic mixer at a volume ratio of 1:3, with a combined flow rate of 50.0 mL/min. The resulting LNP suspension was dialyzed and concentrated to approximately 0.2 mg/mL using Amicon Ultra centrifugal filters (100-kDa cutoff; Merck) and then filtered through a 0.22-µm filter (Sartorius, Aubagne, France). The Z-average particle size, polydispersity index (PDI), and zeta potential were measured using a Zetasizer Ultra (Malvern Panalytical, Malvern, United Kingdom) (Table S1). The concentrations of total and encapsulated IVT mRNA were quantified using the Quant-it RiboGreen RNA Assay Kit (Thermo Fisher Scientific) in the presence and absence of Triton X-100.

### Mice

Specific pathogen-free male CBA/N mice (Japan SLC, Hamamatsu, Shizuoka, Japan) were housed under a 12-h light/12-h dark cycle with free access to food and water. All mouse experiments were approved by the Animal Care and Use Committee at the Research Institute for Microbial Diseases, the University of Osaka and conducted in accordance with the institutional guidelines of The University of Osaka for the ethical treatment of animals.

### Guinea pigs

Female Hartley guinea pigs (Charles River Laboratories, Wilmington, MA, USA) were housed in animal biosafety level 2 and 4 facilities in the Galveston National Laboratory (GNL) at The University of Texas Medical Branch (UTMB). Guinea pig studies were reviewed and approved by the Institutional Animal Care and Use Committee at UTMB and conducted in accordance with the National Institutes of Health guidelines. All procedures were performed in accordance with institutional ethical guidelines and the American Veterinary Medical Association (AVMA) Guidelines for the Euthanasia of Animals (2020 Edition).

### Viruses

To generate rML29, BHK-21 cells were seeded in a six-well plate, cultured overnight, then transfected with two plasmids based on the reference sequences (NCBI accession nos. NC_006573.1 and NC_006572.1) that direct polymerase I (pol-I)-mediated intracellular synthesis of ML29 S and L RNAs in antigenomic polarity, together with pC-ML29-NP and pC-ML29-L, which drive polymerase II (pol-II)-mediated expression of the ML29-derived trans acting factors NP and L, using Lipofectamine 2000 (Thermo Fisher Scientific). Then, they were incubated at 37°C in 5% CO_2_. Five hours after transfection, the transfection mixture was replaced with fresh medium. Following a 3-day incubation at 37°C in 5% CO_2_, the cell culture medium (cell culture supernatant, CCS) was removed, fresh medium was added, and the plates were incubated at 37°C in 5% CO_2_ for an additional 3 days. CCS collected 6 days post-transfection was used to amplify the rescued virus, using Vero E6 cells as a substrate. The rescue of rLCMV/LASgpc^2m^ was performed as described (*26*). The LASV strain LF2384 was propagated in Vero cells as described (*54*). All work with infectious LASV was performed in the BSL-4 facilities of the GNL at UTMB in accordance with institutional guidelines.

### Immunization with mRNA-LNP and rML29 challenge of CBA/N mice

Six-week-old CBA/N mice were i.m. immunized with LASgpc- or LASnp-mRNA-LNP, each containing 2 µg of IVT mRNA, or were mock-immunized with saline, with a 21-day interval between immunizations. Fourteen days after the second immunization, blood samples were collected to evaluate antigen-specific antibody production. Twenty-eight days after the second immunization, mice were inoculated i.c. with 1 × 10^3^ plaque-forming units (PFU) of rML29 and monitored daily for survival.

### Immunization with mRNA-LNP and LASV challenge of guinea pigs

Six-week-old Hartley guinea pigs were i.m. immunized with a bivalent vaccine combining optLASgpc-mRNA-LNP and optLASnp-mRNA-LNP at a high (5 µg mRNA each for the first immunization and 2.5 µg mRNA each for the second immunization) or low (1 µg mRNA each) dose with a 21-day interval between immunizations. Saline was used for mock immunizations. Fourteen days after the second immunization, blood was collected from the vena cava under anesthesia, and serum was separated for antibody tests. Twenty-eight days after the second immunization, guinea pigs were challenged i.p. with 1,000 PFU of LASV (strain LF2384) and monitored daily for 28 days. Body weights were measured with an Ohaus CX1201 Portable Scale. Body temperature was measured using a subcutaneously implanted BMDS TP-1000 transponder and a DAS-8037 transponder reader (Bio Medic Data Systems).

### Statistics

GraphPad Prism 11 (GraphPad, San Diego, CA, USA) was used for all statistical analyses. Statistically significant differences were determined by the two-tailed Mann–Whitney U test.

Additional materials and methods are described in the SI Materials and Methods.

### List of Supplementary Materials

SI Materials and Methods

Figs. S1 to S5

Table S1

## Acknowledgments

We thank Xiuyuan Lu, Mai Hayasaka, Keiko Shindo, and Ayako Takashima for their technical advice and assistance. We thank J. Ludovic Croxford, PhD, from Edanz (https://jp.edanz.com/ac) for editing a draft of this manuscript.

## Author contributions

Conceptualization: MH, MI Methodology: MH, TS, TM, JM, MI

Investigation: MH, TS, RN, TM, SY, GS, DK, CC, KL, ES, JM

Visualization: MH

Funding acquisition: TM, JM, MI Project administration: MH, MI Resources: TM, ISL, JCT, JM, MI Supervision: MI

Writing – original draft: MH, MI

Writing – review & editing: MH, TS, RN, TM, DK, ISL, JCT, JM, MI

## Data and materials availability

All data are available in the main text or in the supplementary materials.

