## Supplementary Information for "A bivalent Lassa virus mRNA vaccine protects guinea pigs from lethal infection with Lassa virus"

### SI Materials and Methods

#### Cells

Vero E6 (CRL-1586; American Type Culture Collection, ATCC, Manassas, VA, USA), Vero (CRL-81; ATCC), and HEK293 (CRL-1573; ATCC) cells were maintained in Dulbecco's modified Eagle's medium (DMEM) supplemented with 10% heat-inactivated fetal bovine serum (FBS), 100 U/mL penicillin, and 100 μg/mL streptomycin (1% P/S) (10% FBS/DMEM) at 37°C in 5% $CO_2$. BHK-21 (CRL-3216; ATCC) cells were grown in 10% FBS/DMEM with an additional 5% tryptose phosphate broth under the same conditions of 37°C and 5% $CO_2$.

#### Virus titration

rML29 titers were determined by a standard plaque assay. Briefly, confluent monolayers of Vero E6 cells in 6-well plates were inoculated with serial dilutions of rML29. After incubation at 37°C in 5% $CO_2$ for 60 min, virus inoculum was removed and a semi-solid solution consisting of 0.5% Seakem ME (Lonza, Rockland, ME, USA) in modified Eagle's medium (MEM; Thermo Fisher Scientific, Waltham, MA, USA) supplemented with 0.7% FBS and 1% P/S was added to each well. The plates were then incubated at 37°C in 5% $CO_2$. After 5 days of incubation, cells were fixed with 10% formalin, and plaques were visualized by crystal violet staining. The number of plaques was counted to calculate the viral titer. rLCMV/LASgpc$^{2m}$ titers were determined by an immunofocus forming assay as described previously (*1*). Lassa virus (LASV; strain LF2384) titers were determined by plaque assay as described previously (*2*).

#### Assessment of the LASV glycoprotein precursor (GPC) and nucleoprotein (NP) expression in cells transfected with i*n vitro*-transcribed (IVT) mRNAs

HEK293 cells seeded in 12-well plates at 4.5 × $10^5$ cells per well and cultured overnight were transfected with 1.8 μg of *in vitro*-transcribed (IVT) mRNAs or mock-transfected using 4.5 μL of Lipofectamine 2000 (Thermo Fisher Scientific) and incubated at 37°C in 5% $CO_2$. At 5 h post-transfection, the transfection mixture was removed, fresh medium was added to the wells, and the cells were incubated at 37°C in 5% $CO_2$. At 24 h post-transfection, total cell lysates were prepared by lysing the cells with PD buffer (0.5% Triton X-100, 250 mM NaCl, 50 mM Tris-HCl, 10% glycerol, 1 mM $MgCl_2$, 1 μM $CaCl_2$) containing Halt Protease and Phosphatase Inhibitor Cocktail (Thermo Fisher Scientific). The resulting cell lysate was clarified by centrifugation at 20,600 × *g* and 4°C for 5 min. The clarified cell lysate was mixed at a 3:1 ratio with 4× LDS sample buffer (Thermo Fisher

Scientific) containing 10× sample reducing agent (Thermo Fisher Scientific) and denatured at 70°C for 10 min. Protein samples were fractionated by SDS-polyacrylamide gel electrophoresis using a 4%–12% Bis-Tris Plus Mini Protein Gel (Thermo Fisher Scientific), and the resolved proteins were transferred by electroblotting onto polyvinylidene difluoride membranes (Immobilon-P PVDF Transfer Membranes; Merck, Darmstadt, Germany). To detect LASV GPC, NP, or GAPDH, the membranes were incubated with a rabbit polyclonal antibody to LASV GP2 (PA5-117438; Thermo Fisher Scientific), a rabbit monoclonal antibody to LASV NP (GTX636830; GeneTex, Irvine, CA, USA), or a mouse monoclonal antibody to GAPDH (M171-3; MBL, Tokyo, Japan) and then with horseradish peroxidase (HRP)-conjugated anti-rabbit IgG or anti-mouse antibodies (Jackson ImmunoResearch Laboratories, West Grove, PA, USA), as appropriate. The Chemi-Lumi One L chemiluminescent substrate (Nacalai Tesque, Kyoto, Japan) was used to generate chemiluminescent signals that were visualized with a chemiluminescent imager (Amersham ImageQuant 800; Cytiva, Tokyo, Japan).

**Detection of LASV GPC- and NP-specific antibodies in CBA/N mice plasma by ELISA**

Recombinant LASV GPC (CSB-CF362480LNP; CUSABIO, Wuhan, Hubei, China) or NP (CSB-EP318401LNPE1; CUSABIO) was diluted with ELISA Coating buffer (Abcam, Cambridge, UK), coated onto ELISA plates (BioLegend, San Diego, CA, USA) at 20 ng/well, and incubated at 4°C overnight. Plates were washed six times with phosphate-buffered saline (PBS)-T [PBS supplemented with 0.2% polyoxyethylene sorbitan monolaurate (Nacalai Tesque)] before 300 μL of blocking buffer [PBS-T containing 3% normal goat serum (Nacalai Tesque) and 2% skim milk (Nacalai Tesque)] were added and incubated at 37°C for 2 h. Two-fold serial dilutions of plasma were added to the plates, followed by overnight incubation at 4°C. Anti-LASV GP2 polyclonal antibody (PA5-117438; Thermo Fisher Scientific) and anti-LASV NP antibody (GTX134873; GeneTex) were used as positive controls for the LASV GPC- and NP-coated plates, respectively. The plates were washed six times with PBS-T before the addition of anti-mouse IgG-HRP (Jackson ImmunoResearch Laboratories) and incubated at 37°C for 1 h. The plates were washed again with PBS-T and developed with TMB substrate [ELISA POD Substrate TMB Kit (HYPER), Nacalai Tesque]. The reaction was stopped by the addition of stop solution, and the optical density was measured at 450 nm ($OD_{450}$) using a multi-mode plate reader (SpectraMax iD5, Molecular Devices, San Jose, CA, USA).

**Focus reduction neutralizing test (FRNT)**

FRNT was performed as previously described (*1*). Briefly, confluent monolayers of Vero E6 cells were inoculated at a multiplicity of infection of 0.05 with a recombinant lymphocytic choriomeningitis virus (LCMV) expressing LASV GPC (rLCMV/LASgpc$^{2m}$) preincubated with two-fold serial dilutions of mouse plasma at 37°C for 1 h. A murinized neutralizing antibody (nAb) clone 37.7H (Ab01102-2.0; Absolute Antibody, Redcar, Cleveland, UK) was used as a positive control. At 20 h post-inoculation, the cells were fixed with 4% paraformaldehyde in PBS (4% PFA/PBS) (Nacalai Tesque). After cell permeabilization and blocking with dilution buffer [0.3% Triton X-100 (Sigma-Aldrich, St. Louis, MO, USA) in PBS containing 3% bovine serum albumin (Nacalai Tesque)], the cells were incubated with a monoclonal antibody against LCMV NP (VL-4, Bio X Cell, Lebanon, NH, USA) conjugated with Alexa Fluor 488 (Alexa Fluor 488 Protein Labelling Kit; Thermo Fisher Scientific). Fluorescent images were acquired using a CQ1 Confocal Quantitative Image Cytometer (Yokogawa Electric Corporation, Tokyo, Japan), and the NP-positive LCMV-focus numbers were determined using the high-content analysis software CellPathfinder (Yokogawa Electric Corporation).

**LASV GPC immunodominant T-cell epitope mapping**

For LASV GPC (lineage IV, Josiah strain) CD8$^+$ T-cell epitope mapping, 57 9-mer peptides spanning aa positions 57–83, 218–244, and 400–426 of LASV GPC with a 1-aa offset were individually synthesized (GenScript, Piscataway, NJ, USA) and dissolved in DMSO. Eight-week-old CBA/N mice were inoculated intraperitoneally (i.p.) with 1 × $10^3$ PFU of rML29. At 8 days post-inoculation (dpi), spleens were collected from euthanized mice and erythrocyte-free splenocytes were prepared. LASV GPC-specific T-cell responses were assessed using a mouse IFN-γ ELISPOT assay kit (Cellular Technology Ltd., Cleveland, OH, USA). Briefly, erythrocyte-free splenocytes were added to a 96-well filter plate pre-coated with anti-mouse monoclonal IFN-γ antibody at 3 × $10^5$ cells per well and stimulated overnight at 37°C in 5% $CO_2$ with 2 µg/mL of each 9-mer peptide. Cell stimulation cocktail (Thermo Fisher Scientific) and CTL medium alone were used as positive and negative controls. After stimulation, plates were developed according to the manufacturer's instructions. Spot-forming cells (SFC) were counted using ImmunoSpot S6 VERSA Analyzer (Cellular Technology Ltd.). Further fine-mapping was performed using the mouse IFN-γ ELISPOT assay with four 8–10-mer peptides (GenScript) spanning aa 65–73 of LASV GPC (Fig. 2B). For LASV GPC-specific CD4$^+$ T-cell mapping, 20- or 15-mer peptides covering aa 218–239 (Fig. 2C) were synthesized (GenScript) and used in the mouse IFN-γ ELISPOT assay as described above, with or without the depletion of CD4$^+$ or CD8$^+$ T cells using mouse CD4 (L3T4) or CD8a (Ly-2) MicroBeads, respectively, followed by magnetic separation on LD columns (Miltenyi Biotec, Bergisch Gladbach, Germany).

#### LASV NP immunodominant T-cell epitope mapping

For LASV NP (lineage IV, Josiah strain) T-cell epitope mapping, an overlapping peptide library of 140 15-mer peptides covering the full-length LASV NP sequence with a 4-aa offset was synthesized (GenScript) and dissolved in DMSO. Twenty-four peptide pools (A–X) were generated by mixing 8–12 peptides according to a matrix pooling strategy (Fig. 3A). Antigen-specific IFN-γ induction was examined using the mouse IFN-γ ELISPOT assay described above with the peptide pools. Four peptides located at the intersections of responsive peptide pools were further analyzed for $CD4^+$ or $CD8^+$ responsiveness following the depletion of $CD4^+$ or $CD8^+$ T cells as described above. To identify H-$2K^k$-restricted LASV NP-specific $CD8^+$ T-cell epitopes, we conducted *in silico* screening using 10 $CD8^+$ T-cell epitope prediction algorithms available through the Immune Epitope Database T Cell Epitope Prediction (https://www.iedb.org/home_v3.php). The top three peptides with the highest scores were selected from each algorithm. After the removal of duplicates, 16 peptides were synthesized (GenScript), dissolved in DMSO, and experimentally evaluated using the mouse IFN-γ ELISPOT assay described above.

#### Intracellular cytokine staining (ICS)

Eight-week-old CBA/N mice were inoculated i.p. with $1 \times 10^3$ PFU of rML29 or mock-immunized with saline. At 8 dpi, the spleens were harvested from euthanized mice, and erythrocyte-free splenocytes were prepared. Alternatively, 6-week-old CBA/N mice were immunized i.m. twice with mRNA-LNP containing 2 µg of IVT mRNA, or mock-immunized with saline, at an interval of 21 days. Eight days after the second immunization, spleens were harvested from euthanized mice, and erythrocyte-free splenocytes were prepared. The resulting splenocytes ($2 \times 10^6$ cells) were stimulated with 2 µg/mL of the peptides specified in the corresponding figures, and 50 U/mL of recombinant mouse IL-2 (BioLegend) overnight, followed by a 4-hour incubation with Protein Transport Inhibitor Cocktail (Thermo Fisher Scientific). As a positive control, Cell Stimulation Cocktail (Thermo Fisher Scientific) was used instead of the peptides. Cells were stained with cell surface marker antibodies (CD3ε: PerCP/Cyanine5.5, clone 145-2C11; CD8a: FITC, clone 53-6.7; CD4: Pacific Blue, clone RM4-5, all from BioLegend) and Fixable Viability Dye eFluor 780 (Thermo Fisher Scientific). Following cell fixation with 4% PFA/PBS and permeabilization with Perm/Wash Buffer (BD Biosciences, San Jose, CA, USA), intracellular IFN-γ staining was performed on half of the cells using an anti-IFN-γ antibody

(PE, clone XMG1.2; BioLegend), and on the other half using an isotype control antibody (rat IgG1 κ: PE, clone RTK2071; BioLegend). Flow cytometric analysis was performed using an SH800H flow cytometer (Sony, Tokyo, Japan), and the data were analyzed using Cell Sorter Software (Sony).

**Detection of LASV GPC- and NP-specific antibodies in guinea pig serum by ELISA**

To detect LASV GPC- or NP-specific antibodies in guinea pig serum, ELISAs were performed as previously described (*3, 4*). For anti-LASV GPC antibody detection, purified LASV glycoprotein was used as an antigen (*4*). For anti-LASV NP antibody detection, whole cell lysate overexpressing LASV NP was used as an antigen (*3*). Antibodies bound to the antigens were detected by goat Anti-Guinea pig IgG H&L (HRP) (Abcam) and visualized by adding 3,3',5,5'-tetramethylbenzidine Liquid Substrate, Supersensitive, for ELISA (Sigma-Aldrich) and the reaction was stopped with 1M phosphoric acid. The $OD_{450}$ was measured and normalized to the $OD_{450}$ of negative control antigens. For the anti-LASV GPC ELISA, purified recombinant EBOV GP-His (*5*) was used as a negative control antigen. For the anti-LASV NP ELISA, whole cell lysate from cells transfected with the empty pCAGGS vector was used.

**Plaque reduction neutralizing test (PRNT)**

PRNT was performed as described previously (*6*). Briefly, LASV strain LF2384 (500 PFU per well) was preincubated with heat-inactivated guinea pig serum diluted 1:10 using DMEM containing 2% FBS at 37°C for 30 min and inoculated onto confluent monolayers of Vero cells in 6-well plates. After incubation at 37°C for 30 min, the virus inoculum was removed, and MEM containing 0.6% Tragacanth (Sigma-Aldrich) and 2% FBS was added to each well. After 5 days of incubation, cells were fixed with 10% formalin, and plaques were visualized by crystal violet staining. The number of plaques was counted manually using a light box.

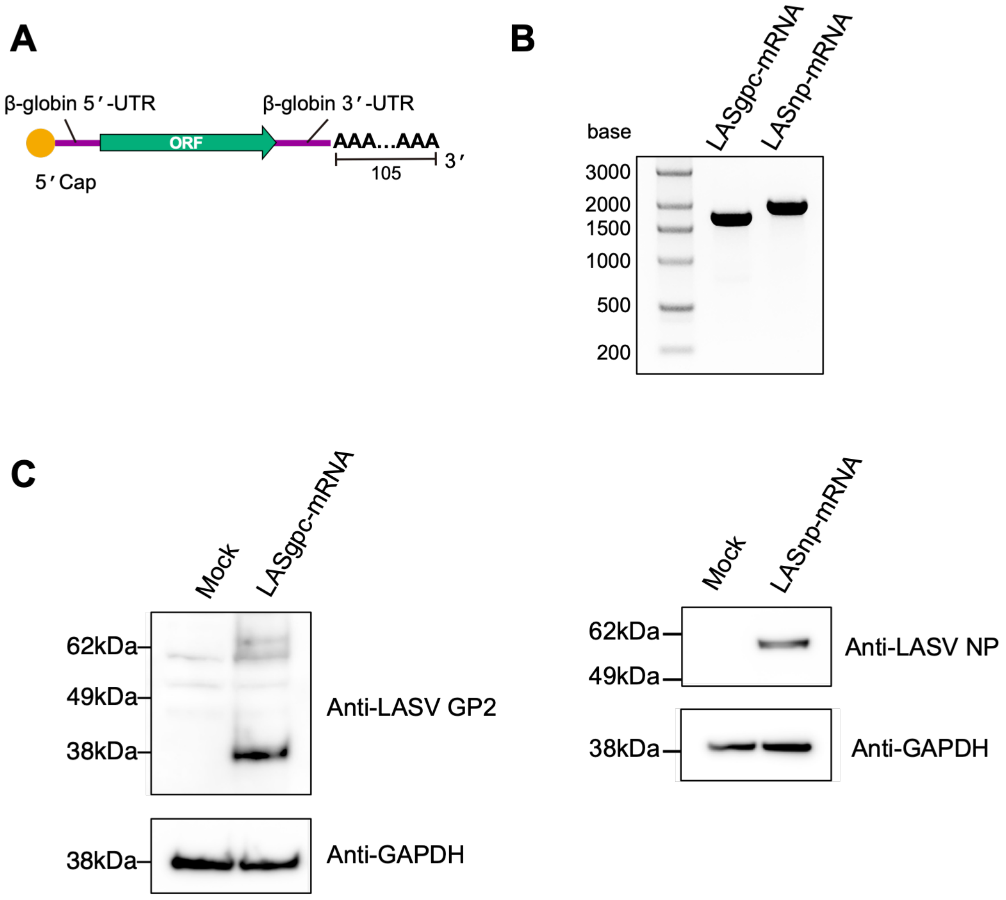


**Fig. S1. Generation of IVT mRNAs encoding LASV GPC or NP.** (**A**) Schematic diagram of the IVT mRNA constructs. mRNAs containing 5′ cap 1 structure, human β-globin 5′ UTR, an open reading frame of LASV GPC (LASgpc-mRNA) or NP (LASnp-mRNA), human β-globin 3′ UTR, and a 105-nt poly(A) tail were transcribed *in vitro* from linearized DNA templates. (**B**) Agarose gel electrophoresis of IVT mRNAs (**C**) Western blot analysis of LASV GPC (left) and NP (right) expression in lysates of HEK293 cells transfected with the corresponding IVT mRNAs.

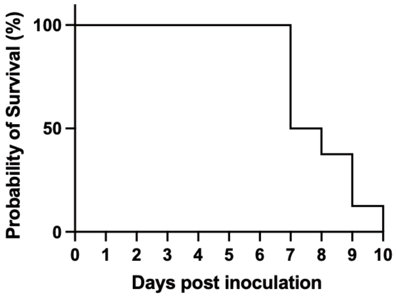


**Fig. S2. Confirmation of the lethal CBA/N-rML29 model.** Eight-week-old CBA/N mice (n = 8) were inoculated i.c. with 1 × $10^3$ PFU of rML29 and survival was monitored daily.

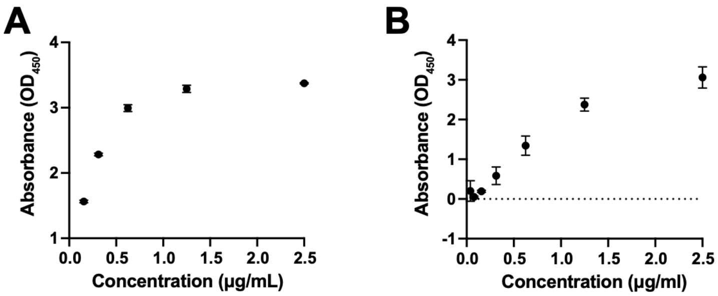


**Fig. S3. Verification of ELISAs used to detect LASV GPC- and NP-specific antibodies in plasma from CBA/N mice.** (**A**, **B**). The optical density at 450 nm ($OD_{450}$) was determined for two-fold serial dilutions of anti-LASV GP2 polyclonal antibody (PA5-117438) (**A**) and anti-LASV NP polyclonal antibody (GTX134873) (**B**) using the corresponding ELISAs. The $OD_{450}$ values from samples without antibody were subtracted as background.

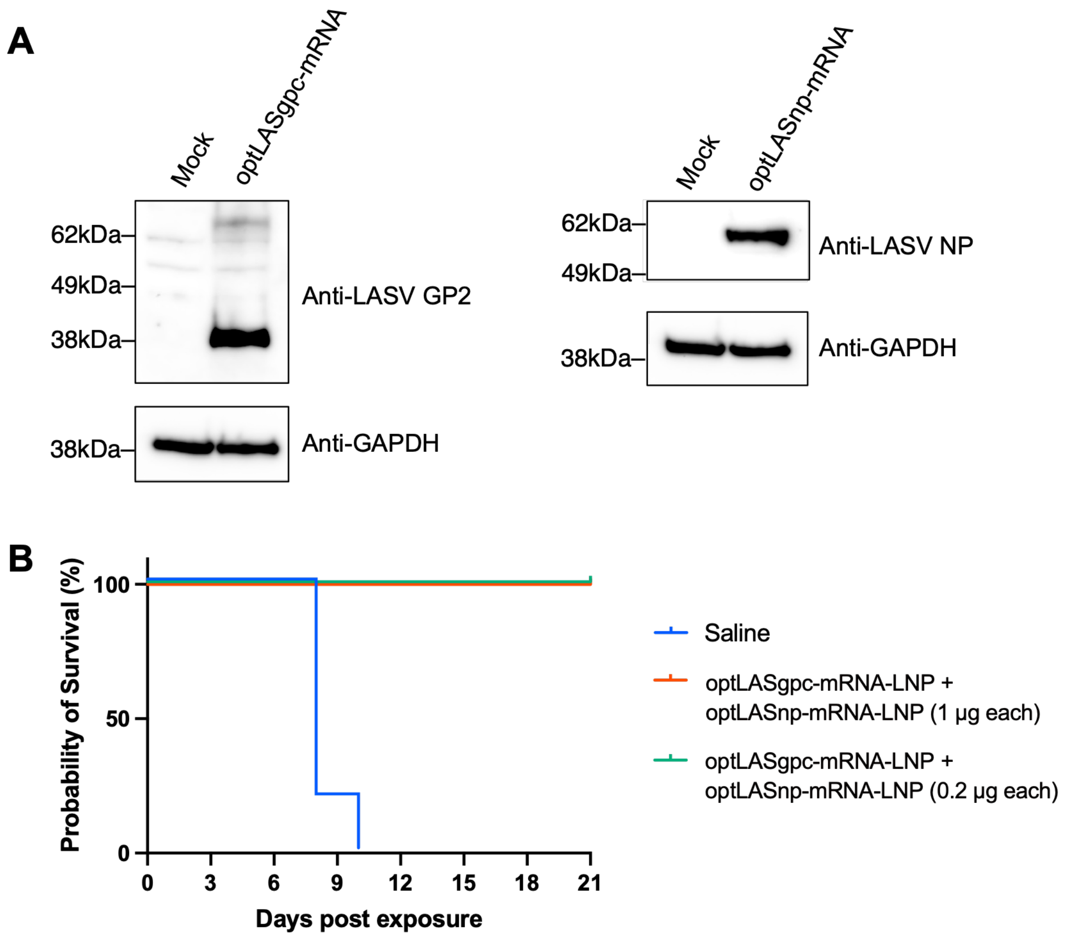


**Fig. S4. Characterization of optLASgpc-mRNA-LNP and optLASnp-mRNA-LNP.** (**A**) Western blot analysis of LASV GPC and NP expression in lysates of HEK293 cells transfected with optLASgpc-mRNA (left) and optLASnp-mRNA (right). (**B**) Survival of CBA/N mice (n = 5 per group) immunized twice, 21 days apart, with a bivalent vaccine combining optLASgpc-mRNA-LNP and optLASnp-mRNA-LNP at a high (1 µg mRNA each) or low (0.2 µg mRNA each) dose, or mock-immunized with saline, following challenge with 1 × $10^3$ PFU of rML29 28 days after the second immunization.

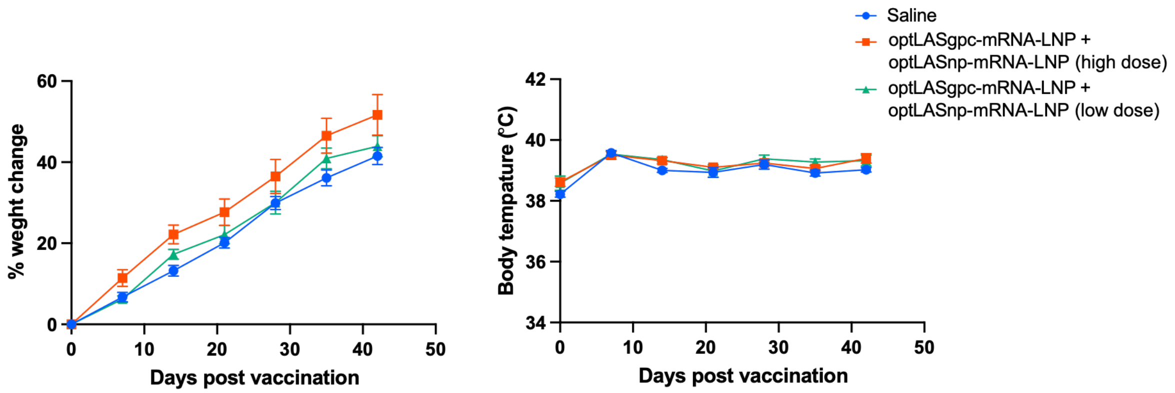


**Fig. S5. Safety evaluation of the bivalent LASV mRNA vaccine.** Body weight (left) and body temperature (right) of guinea pigs immunized twice, 21 days apart, with a bivalent vaccine combining optLASgpc-mRNA-LNP and optLASnp-mRNA-LNP at a high (5 μg mRNA each for the first immunization and 2.5 μg mRNA each for the second immunization) or low (1 μg mRNA each) dose, or mock-immunized with saline. First immunization was set as day 0. Data are shown as means ± SD.

**Table S1. Physicochemical characterization of mRNA-LNPs**

| mRNA-LNP | Z-Average (nm) | PDI | Encapsulation Efficiency (%) | ζ-potential (mV) |
|---|---|---|---|---|
| optLASgpc-mRNA-LNP | 73.88 ± 2.04 | 0.08 ± 0.02 | 96.1 ± 1.65 | -7.73 ± 0.81 |
| optLASnp-mRNA-LNP | 73.76 ± 1.86 | 0.09 ± 0.04 | 95.3 ± 1.30 | -7.70 ± 1.08 |

Data are presented as the mean ± SD of three independent preparations.
